# Genomics analysis of hexanoic acid exposure in *Drosophila species*

**DOI:** 10.1101/2021.06.08.447576

**Authors:** Z.A. Drum, S.M. Lanno, S.M. Gregory, S.J. Shimshak, M. Ahamed, W. Barr, B. Bekele, A. Biester, C. Castro, L. Connolly, N. DelGaudio, W. Humphrey, H. Karimi, S. Karolczak, T. Lawrence, A. McCracken, N. Miller-Medzon, L. Murphy, C. Park, S. Park, C. Qiu, K. Serra, G. Snyder, A. Strauss, S. Tang, C. Vyzas, Joseph D. Coolon

## Abstract

*Drosophila sechellia* is a dietary specialist endemic to the Seychelles islands that has evolved to consume the fruit of *Morinda citrifolia*. When ripe, the fruit of *M. citrifolia* contains octanoic acid and hexanoic acid, two medium chain fatty acid volatiles that deter and are toxic to generalist insects. *D. sechellia* has evolved resistance to these volatiles allowing it to feed almost exclusively on this host plant. The genetic basis of octanoic acid resistance has been the focus of multiple recent studies, but the mechanisms that govern hexanoic acid resistance in *D. sechellia* remain unknown. To understand how *D. sechellia* has evolved to specialize on *M. citrifolia* fruit and avoid the toxic effects of hexanoic acid, we exposed adult *D. sechellia, D. melanogaster* and *D. simulans* to hexanoic acid and performed RNA sequencing comparing their transcriptional responses to identify *D. sechellia* specific responses. Our analysis identified many more genes responding transcriptionally to hexanoic acid in the susceptible generalist species than in the specialist *D. sechellia*. Interrogation of the sets of differentially expressed genes showed that generalists regulated the expression of many genes involved in metabolism and detoxification whereas the specialist primarily downregulated genes involved in the innate immunity. Using these data we have identified interesting candidate genes that may be critically important in aspects of adaptation to their food source that contains high concentrations of HA. Understanding how gene expression evolves during dietary specialization is crucial for our understanding of how ecological communities are built and how evolution shapes trophic interactions.

## Introduction

Insects have long been recognized as one of the most abundant and diverse groups of organisms on the planet, with a large fraction of them feeding on plants (Jaenike *et al*. 1990, Stork 2018). Many of these phytophagous insects have evolved to be highly host plant specific. The evolution of such specialized interactions is often guided by specific plant chemistry, with most plants responding to increased insect herbivory by the production of toxic secondary metabolites (Jaenike *et al*. 1990, Petschenka and Agrawal 2016). This leads to an evolutionary arms race as insects evolve resistance to these toxins (Heidel-Fischer and Vogel 2015). While host plant-insect adaptions are well studied in the literature, less is known about the underlying genetic mechanisms that contribute to the evolution of these complex ecological interactions (Ungerer *et al*. 2007).

*Drosophila sechellia* feeds and oviposits primarily on *Morinda citrifolia*, a fruit highly toxic to other *Drosophila* species (Legal *et al*. 1992). The plant produces ripe fruit year-round in the Seychelles island archipelago, the sole location where *D. sechellia* are found (Legal *et al*. 1992) providing abundant and consistent resources. Upon exposure to the ripe fruit, other *Drosophila* species display frantic behavior and wing movements, reduction in locomotor activity, and death (Legal *et al*. 1994). *Drosophila sechellia* eggs are able to hatch and develop on *M. citrifolia* fruit but the embryos of other *Drosophila* species die (Amlou *et al*. 1998). Unlike its generalist sister species, *D. sechellia* prefer *M. citrifolia* to other hosts and are drawn to the fruit from a long distance (R’Kha *et al*. 1991). *Drosophila sechellia* evolved resistance to the toxins in *M. citrifolia* from an ancestral sensitive state (R’Kha *et al*. 1997) and this tolerance provides *D. sechellia* a temporal advantage over other fruit fly species that can only lay eggs in the fruit once it has rotten and the toxic volatiles are reduced.

The toxic properties of *Morinda* are attributed to the carboxylic acids present in the pulp of the ripe fruit (Legal *et al*. 1994). This pulp is largely characterized by carboxylic acids, primarily the fatty acids octanoic (OA) and hexanoic acid, (HA) which comprise 58% and 19% of the volatile compounds found the ripe fruit respectively (Farine *et al*. 1996). While the genetic basis of *D. sechellia* resistance to the most abundant and toxic compound in *M. citrifolia* fruit, OA, has been characterized in previous studies (Dworkin and Jones 2009, Andrade Lopez *et al*. 2017, Lanno *et al*. 2017, Peyser *et al*. 2017, Lanno and Coolon 2019, Lanno *et al*. 2019a, Lanno *et al*. 2019b), much less is known about the genes involved in HA resistance. In a study using fatty acid concentrations equivalent to 1.5g of ripe *M. citrifolia* fruit, OA treatment alone killed all fruit fly species assayed except *D. sechellia*, while HA alone caused reversible knock-down in other Drosophilids (Farine *et al*. 1996). In a more recent study using higher concentrations of HA, mortality was observed in response to exposure to HA in *D. melanogaster* and *D. simulans* and *D. sechellia* is significantly more resistant to HA induced mortality than sister species (Peyser et al 2017, Lanno and Coolon 2019). Surprisingly, tests of the three major detoxification gene families, cytochrome P450s (cyps), glutathione-S-transferases (GSTs) and esterases (Ests) found that none was involved in derived HA resistance in *D. sechellia* suggesting an alternative genetic mechanism must be involved in resistance. Studies have suggested that HA may be a more efficient *D. sechellia* attractant than OA, while OA is a more potent repellant of generalist species (Amlou *et al*. 1998). In a test using laboratory food medium supplemented with 0.5% of either OA or HA, *D. sechellia* exhibited oviposition preference for media supplemented with HA over OA (Amlou *et al*. 1998).

From a study analyzing the transcriptomic response of *D. sechellia* on OA, 104 genes were found to be differentially expressed in response to OA (Lanno *et al*. 2017). This study showed that several *Osiris* genes, including *Osi6* are upregulated in *D. sechellia* in response to OA. Another study showed that RNAi mediated knockdown of *Osi6* expression drastically decreased survival in response to OA (Andrade Lopez et al 2017). Given that HA makes up about of fifth of the volatile compounds found in *M. citrifolia* and produces unique effects on behavior (Farine *et al*. 1996) and has an unknown and less common basis for toxin resistance (Lanno and Coolon 2019, Peyser et al 2017), identifying the genes responding to HA is necessary to understand how *D. sechellia* has specialized on *M. citrifolia* fruit, and may help pinpoint genes that are involved in resistance to HA. By also analyzing the genes responding to HA exposure in generalist *Drosophila* species D. melanogaster and D. simulans we can identify derived gene expression responses specific to *D. sechellia* that may be critical for HA associated traits (Coolon et al 2009). In this study, adult female *D. sechellia, D. melanogaster*, and *D. simulans* flies were fed either control food or food supplemented with 0.23% HA and significantly differentially expressed genes (DEGs) were identified using RNA-seq. Comparison of the identified genes with those found to respond to OA (Lanno *et al*. 2017) and L-DOPA (Lanno et al 2019), another highly abundant compound found in *M. citrifolia* fruit, identified several genes common in response to OA, HA, and L-DOPA as well as genes unique to HA suggesting these genes may play an important role in the evolved resistance and specialization of *D. sechellia* to *M. citrifolia*.

## Methods

### Fly strains and culture

*Drosophila sechellia* (14021-0428.25), *D. simulans* (14021-0251.195), and *D. melanogaster* (14021-0231.36) flies were reared on standard cornmeal medium under a 16:8 light:dark cycle maintained at 20°C.

### RNA extraction, library preparation, and sequencing

Zero to three day post eclosion adult female flies were fed control food (0.75g Drosophila instant medium Formula 4-24, Carolina Biological Supply Company) or identical food containing 0.23% hexanoic acid (HA). After 24 hours, three replicates of ten whole flies per species and per treatment were homogenized and total RNA was extracted with a modified protocol of the Promega SV extraction system (Coolon *et al*. 2013, **Figure 1**). RNA quality and quantity was assessed using agarose gel electrophoresis and Nanodrop spectrophotometry. RNA was sent to the University of Michigan Sequencing Core Facility for poly-A selection, cDNA synthesis, bar-coded library preparation with TruSeq library preparation kits and sequencing on an Illumina Hiseq 4000, generating 405,166,795 single-end 65 nt sequencing reads for *D. sechellia* and 51nt sequencing reads for *D. melanogaster* and *D. simulans*. (**Table 1**).

**Figure 1:**
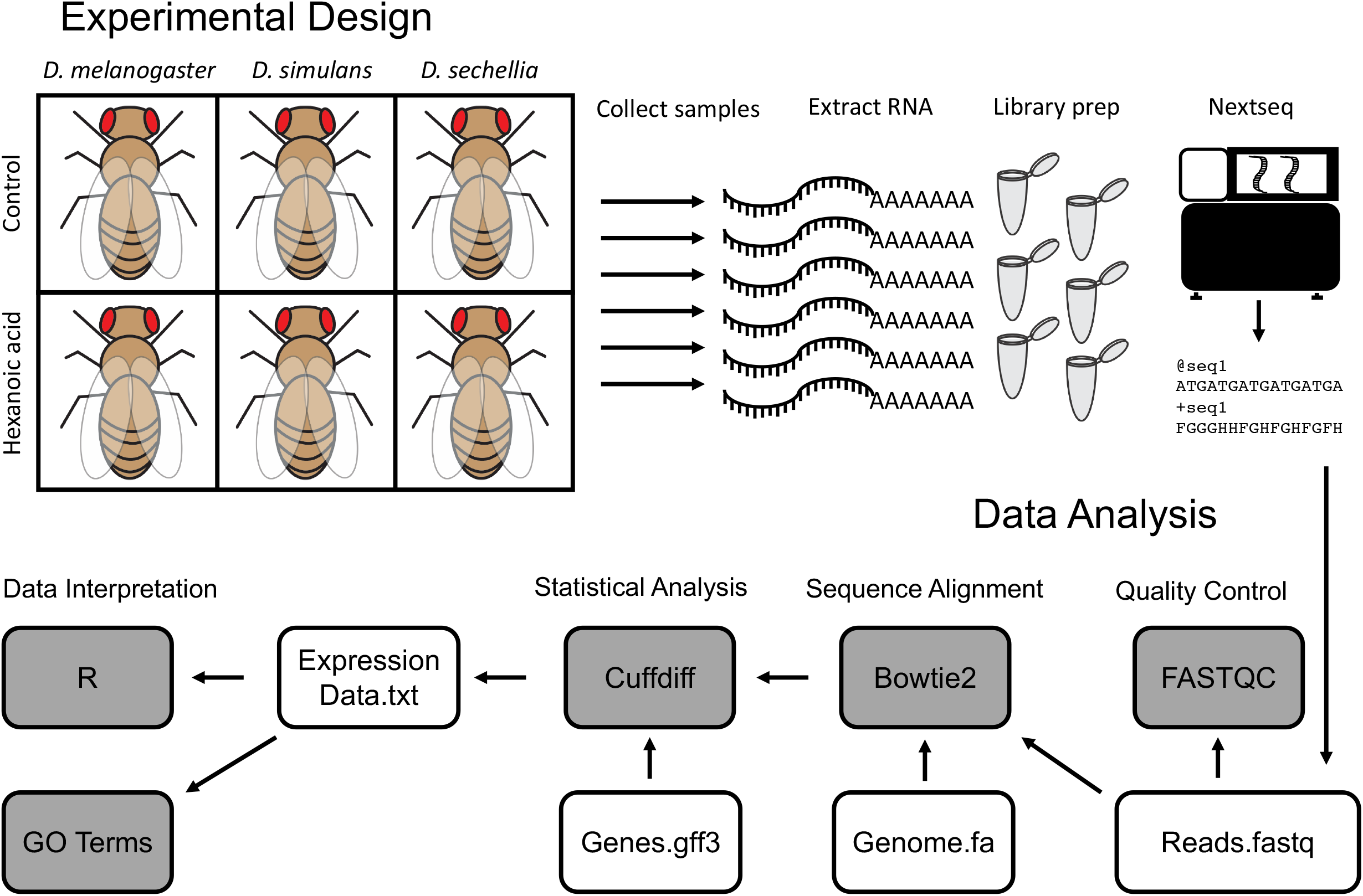
Experimental design and RNA-sequencing pipeline analysis. (A) 0-4 day old adult female *D. sechellia, D. simulans*, and *D. melanogaster* flies were treated for 24 hours on either control food or food supplemented with 0.23% hexanoic acid (HA). Flies were frozen in liquid nitrogen, RNA was extracted, library prep was performed using poly-A selection, and libraries were sequenced. (B) Raw sequencing reads were analyzed using FASTQC for quality control, and then aligned to the respective reference genome using Bowtie2. Differential expression was quantified using Cuffdiff using available respective genome annotation. Data was visualized using R, and significantly differentially expressed genes (GRNs) were used for downstream analysis.

**Table 1:**
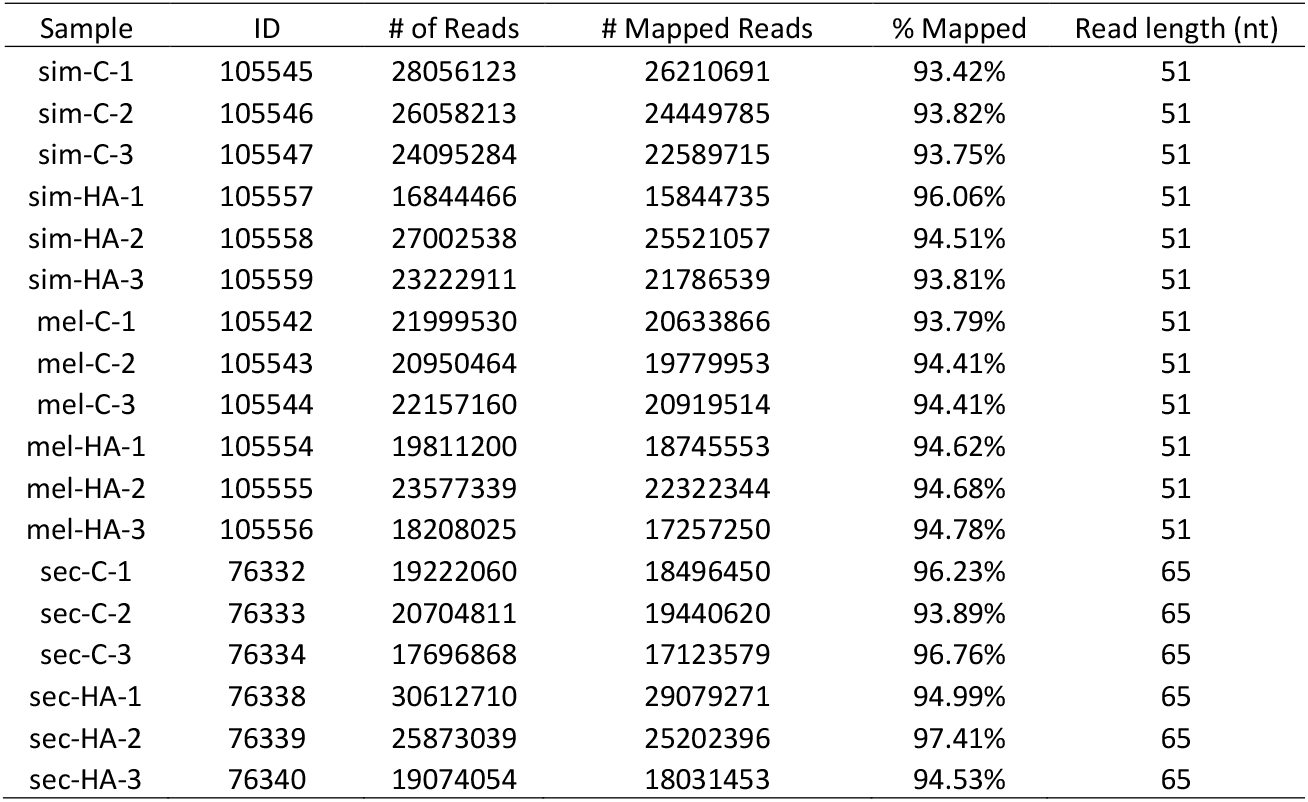
Sequencing results and mapping percentage.

### BIOL310 Genomics Analysis

The genomics analysis of RNA-seq data presented in this manuscript was performed by 20 undergraduate and 6 graduate students as part of a semester-long course at Wesleyan University called Genomics Analysis (BIOL310). This is the third such manuscript (see Lanno *et al*. 2017 and Lanno *et al*. 2019a) made from this course where the aim is to provide undergraduate students an early opportunity with a course-based research experience with active participation in scientific discovery. Students in the course learn through engaging with never-before analyzed data using cutting edge genomics analysis techniques and bioinformatics tools through a discovery-based independent study. Every student in the course contributed to the quality control, analyses, write-up and interpretation of the findings, providing their own unique perspective of the results and text written by each and every student was combined into this manuscript with very little modification.

After sequencing output files were obtained from the University of Michigan Sequencing Core (**Table 1**), fastq files containing raw sequencing reads were uploaded to the Galaxy platform (Afgan *et al*. 2016) and an RNA-seq analysis pipeline was performed (**Figure 1**) as previously described (Lanno *et al*. 2017 and Lanno *et al*. 2019a). Briefly, reads were assessed for quality using FASTQC (Andrews 2010) and any overrepresented sequences were analyzed using NCBI Blast (Altschul *et al*. 1990). Bowtie2 was used for mapping reads to the appropriate reference genome for each species with default parameters (Langmead and Salzberg 2012), with the most recent genomes for each species available at the time of analysis acquired from Ensembl (www.ensembl.org, Yates *et al*. 2016) (*D. sechellia*: Drosophila_sechellia.dsec_caf1.dna.toplevel.fa, *D. simulans*: Drosophila_simulans.ASM75419v3.dna.toplevel.fa and *D. melanogaster*: Drosophila_melanogaster.BDGP6.dna.toplevel.fa). The Bowtie2 output files were analyzed using Cuffdiff (Trapnell *et al*. 2010), which performs gene expression quantification and differential gene expression analysis using the aforementioned genome file along with the most recent annotated .gff3 file for each genome available at the time of analysis acquired from Ensembl (*D. sechellia*: Drosophila_sechellia.dsec_caf1.42.gff3, *D. simulans*: Drosophila_simulans.ASM75419v3.42.gff3 and *D. melanogaster*: Drosophila_melanogaster.BDGP6.95.gff3). In Cuffdiff, geometric normalization and library size correction was performed, along with bias correction using the reference genome, giving an output of DEGs for each species following false discovery rate multiple testing correction (Benjamini & Hochberg 1995, q < 0.05). Data was visualized using R (R Core Team, 2013). In order to compare gene expression results across species, we obtained all 1:1:1 orthologs from *D. sechellia, D. simulans* and *D. melanogaster* from Flybase (Thurmond *et al*. 2019). DEGs following *D. sechellia* exposure to OA or L-DOPA were downloaded from online databases (Lanno *et al*. 2017, Lanno *et al*. 2019a). GO term enrichment was performed on *D. melanogaster* orthologs for each species using GeneOntology.org (www.geneontology.org, Ashburner *et al*. 2000, Carbon *et al*. 2021, Mi *et al*. 2019). KEGG pathway analysis was performed using the *D. melanogaster* ortholog for each DEG from each species (https://www.kegg.jp/kegg/tool/map_pathway1.html, Kanehisa and Sato 2020).

### Data accessibility

All RNA-seq data generated in this manuscript have been submitted to the NCBI Gene Expression Omnibus under accession number XXXXX (to be available at time of publishing). Supplemental Tables for this manuscript have been uploaded to GSA figshare.

## Results

### Differential gene expression in response to HA treatment

In order to identify candidate genes that are important in *D. sechellia* host specialization and evolved resistance to HA we sought genes that have altered expression levels in response to HA exposure. Previous studies have shown that such environmentally plastic gene regulation can indicate importance of that gene’s function in that environment making identified genes good candidates for *D. sechellia* HA associated traits (Coolon et al 2009, Lanno et al 2017, Lanno et al 2019). To quantify gene expression response to HA we performed RNA-sequencing (RNA-seq) on adult female flies after exposure to control food and compared this to flies fed food containing 0.23% HA. Because many of the transcriptional responses to HA might be non-specific, we measured gene expression responses in *D. sechellia, D. melanogaster*, and *D. simulans* to identify those responses (or loss of response) that are restricted to *D. sechellia* representing changes that might contribute to its unique phenotypes.

Using this approach we identified 841 genes differentially expressed by *D. melanogaster* (**Figure 2A,D; Table S1**), 743 genes were differentially expressed by *D. simulans* (**Figure 2B,E; Table S2**) and only 93 genes were differentially expressed in *D. sechellia* (**Figure 2 C,F; Table S3**) in response to HA. No significant difference in the number of upregulated genes (50/93) vs downregulated genes (43/93) was observed in *D. sechellia* in response to HA (Binomial Exact Test, p = 0.1066). In *D. simulans*, there was a significant difference in the number of upregulated genes (69/743) compared to the number of downregulated genes (674/743) in response to HA (Binomial Exact Test, p = 2.2e-16, **Figure 2B,E**). In *D. melanogaster*, there was also a significant difference in the number of upregulated genes (171/841) compared to the number of downregulated genes (670/841) in response to HA (Binomial Exact Test, p = 2.2e-16, **Figure 2A,D**). In *D. sechellia*, there were 39 DEGs identified that responded to HA that do not have annotated *D. melanogaster* orthologs 27 of these genes were 5.8S rRNAs, two snoRNAs, and 7 genes of unknown function. Of the 27 5.8S rRNAs, all 27 were upregulated (Binomial Exact Test, p = 7.451e-09). In *D. simulans*, of the 743 DEGs there were annotated *D. melanogaster* orthologs for 673 genes. For the remainder of the analysis, only genes with known *D. melanogaster* orthologs are considered to allow functional interpretations of DEGs, and for all subsequent analyses the *D. melanogaster* ortholog name was used. This filtering resulted in 673 differentially genes in *D. simulans*, 841 in *D. melanogaster* and 54 in *D. sechellia* used in subsequent analyses.

**Figure 2:**
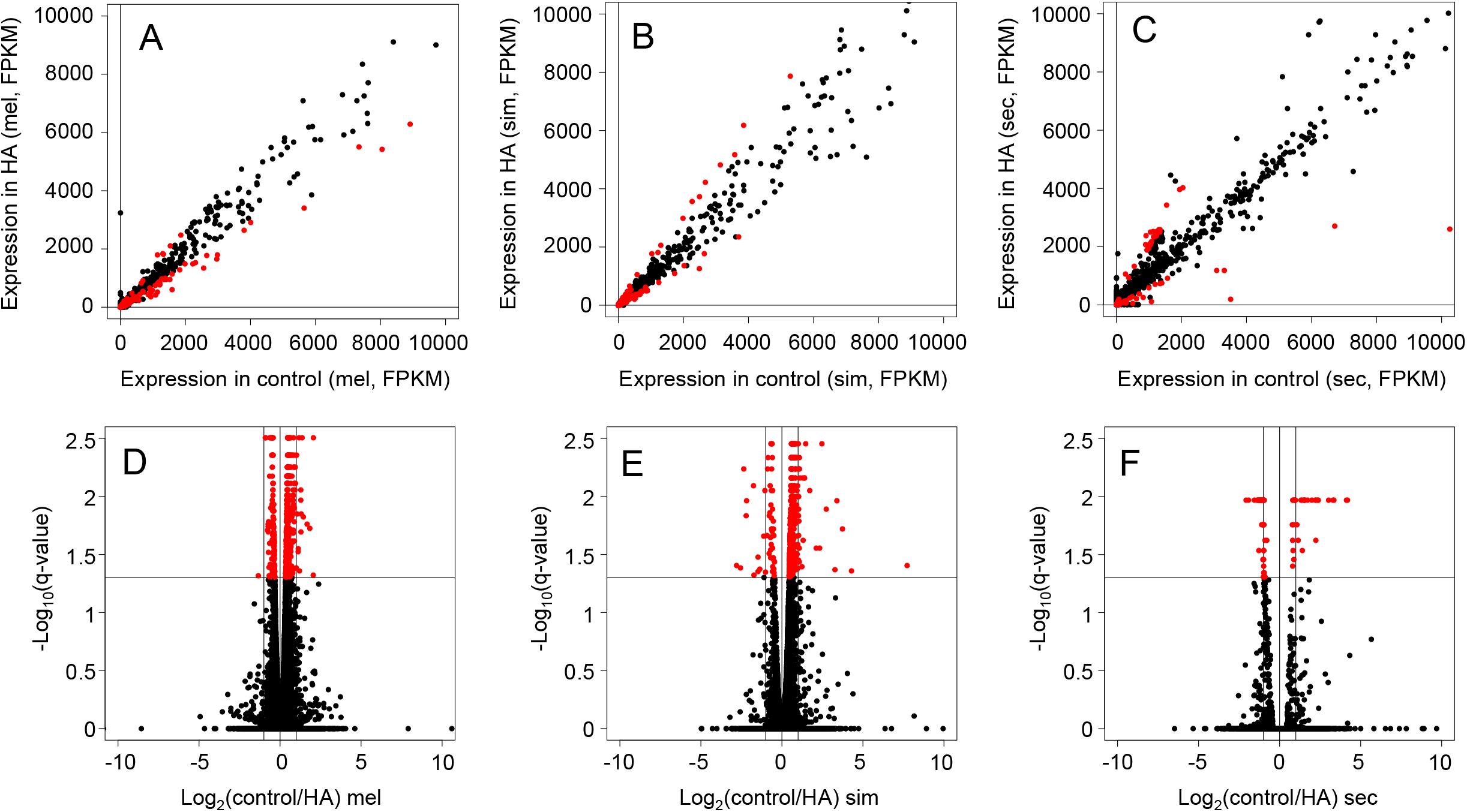
DEGs in adult *D. melanogaster, D. simulans*, and *D. sechellia* in response to HA. (A-C) Plots showing DEG expression in control vs expression in HA treatment in (A) *D. melanogaster*, (B) *D. simulans*, and (C) *D. sechellia*. Statistically significant genes are shown in red. (D-F) Plots showing log_2_(control/HA) fold change in (D) *D. melanogaster*, (E) *D. simulans*, and (F) *D. sechellia*. Statistically significant genes are shown in red.

### Identifying functional enrichment in DEGs

To identify the biological pathways that are involved in responses to HA in *D. sechellia, D. melanogaster*, and *D. simulans* KEGG analyses of the upregulated and downregulated genes in each species were performed. These analyses show that many different metabolic and detoxification pathways along with proteins that localize to the lysosome are changing in response to HA exposure in *D. melanogaster* and *D. simulans*, whereas very few genes in these pathways are responding in *D. sechellia* (**Figures 4A-B**). In *D. sechellia*, many of the of the genes downregulated in response to HA are involved in Toll and Imd immune signaling whereas this was much less prominent in *D. melanogaster* and *D. simulans* DEGs (**Figure 4C**).

To understand which biological and cellular processes are being altered in *D. sechellia, D. melanogaster*, and *D. simulans* in response to HA, Gene Ontology (GO) term enrichment analysis was performed (**Table S16-S18**). In *D. sechellia*, GO term enrichment analysis of DEGs for cellular component showed a significant enrichment for extracellular region genes (p = 4.83e-05), suggesting that several of the genes responding to HA exposure have proteins that are secreted. For biological process GO term enrichment analysis, processes involved in the antibacterial humoral response were significantly enriched (p = 1.75e-08). In *D. sechellia*, no molecular function processes were significantly enriched. In *D. melanogaster*, there was also significant enrichment for GO terms for extracellular region genes (p = 4.42e-04), along with genes found inside the nucleolus (p = 8.40e-04). For biological process, GO terms associated with genes involved in the antibacterial humoral response were significantly enriched as in *D. sechellia* (p = 3.02e-02), as were other processes involved in the *Drosophila* immune response. The *D. melanogaster* HA response also was enriched for genes involved in ribosome biogenesis (p = 3.66e-02). In *D. simulans*, genes found inside the nucleolus were significantly enriched in the set of HA responsive genes (p = 2.65e-02). In an analysis of DEGs in *D. simulans* that are upregulated, genes involved in Notch signaling were significantly enriched (p = 4.11e-04) alongside genes involved in vitelline membrane and chorion formation (p = 2.56e-02). In *D. sechellia*, upregulated genes were significantly enriched for the larval serum protein complex (p = 1.96e-02). Downregulated genes were significantly enriched with antibacterial humoral response GO terms (p = 3.24e-10) along with the response to hyperoxia (p = 1.52e-03). These downregulated genes were also enriched for the extracellular region (p = 4.17e-05). In *D. melanogaster*, upregulated genes were significantly enriched to be involved in the larval serum protein complex (p = 1.23e-03) and were enriched intracellularly (p = 4.02e-16) and within intracellular organelles (1.32e-13). These upregulated genes were significantly enriched in many biological processes, including chromatin silencing (p = 3.25e-03), ecdysone receptor-mediated signaling (p = 1.89e-02), and chorion assembly (p = 2.25e-02). In *D. melanogaster*, downregulated genes were significantly enriched for the antibacterial humoral response (p = 4.52e-03) and for the defense response to Gram-positive bacteria (p = 5.07e-04). These downregulated genes were significantly enriched to be localized to the extracellular region (p = 1.25e-04).

### Comparing DEGs identified in response to HA in D. melanogaster, D. simulans, and D. sechellia

In order to identify genes with *D. sechellia* specific responses to HA, we compared the DEGs from the three species when exposed to HA (**Figure 3A**). We identified 32 genes that were differentially expressed by *D. sechellia* flies exposed to HA that were not responsive to HA exposure in both *D. melanogaster* and *D. simulans* flies (**Figure 3A, Table S13**). In order to identify those genes where *D. sechellia* specific loss of response to HA was observed we selected those genes with significant changes in response to HA in both *D. melanogaster* and *D. simulans* that were not significantly differentially expressed by *D. sechellia* in response to HA. This analysis yielding a total of 213 genes with this expression pattern in our data (**Table S10**). Interestingly, only 2 genes were identified, *CG13114* and *Fbp1*, that had significant response to HA in all three species, with the expression of both increasing in *D. simulans* and decreasing in *D. sechellia* and *D. melanogaster* (**Figure 3A, Table S9**).

**Figure 3:**
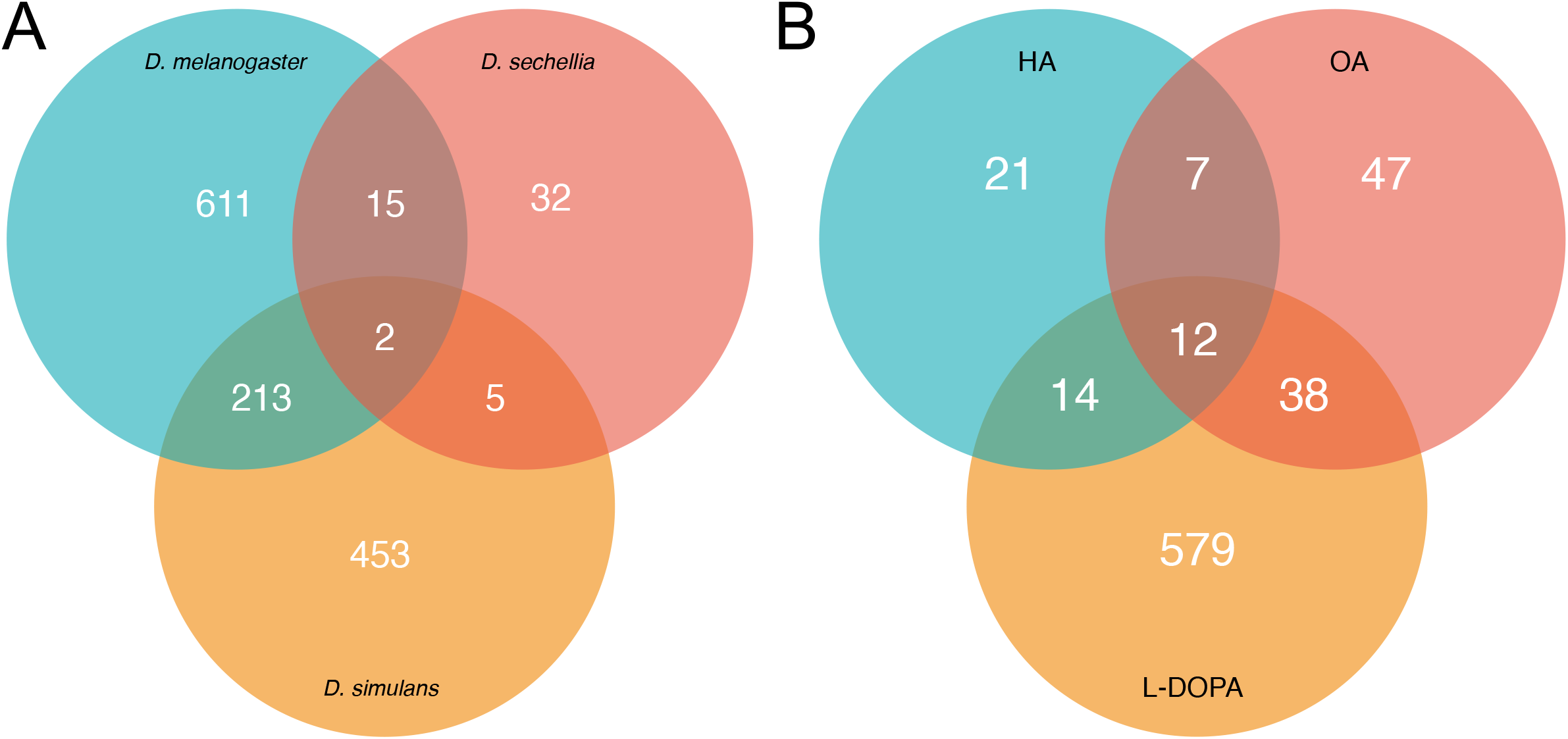
DEGs in *D. melanogaster, D. simulans*, and *D. sechellia* in response to HA. (A) The number of differentially expressed genes in response to HA after RNA-seq are shown for each species, *D. sechellia, D. simulans* and *D. melanogaster*. Overlap and species-specific number of DEGs are indicated. (B) The number of differentially expressed genes in *D. sechellia* when it is exposed to OA, HA, or L-DOPA treatment as well as the number of specific and overlapping genes are shown.

**Figure 4.**
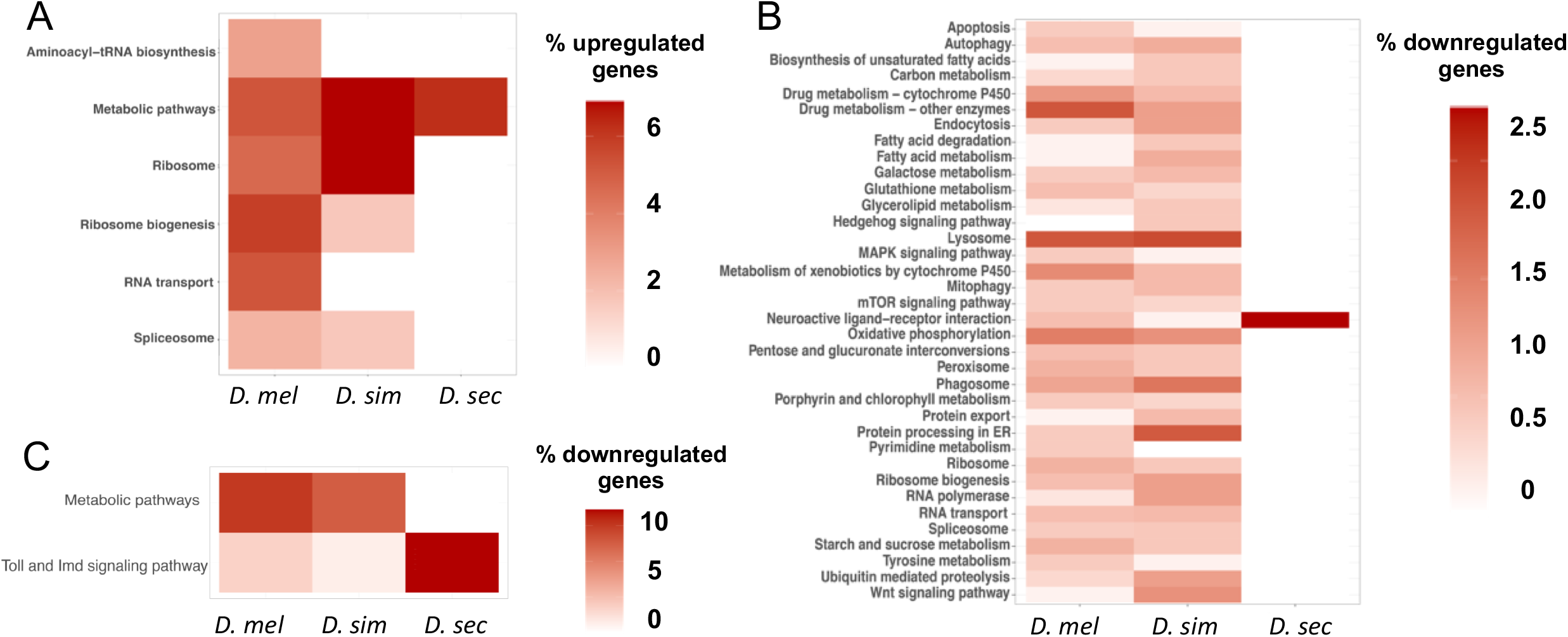
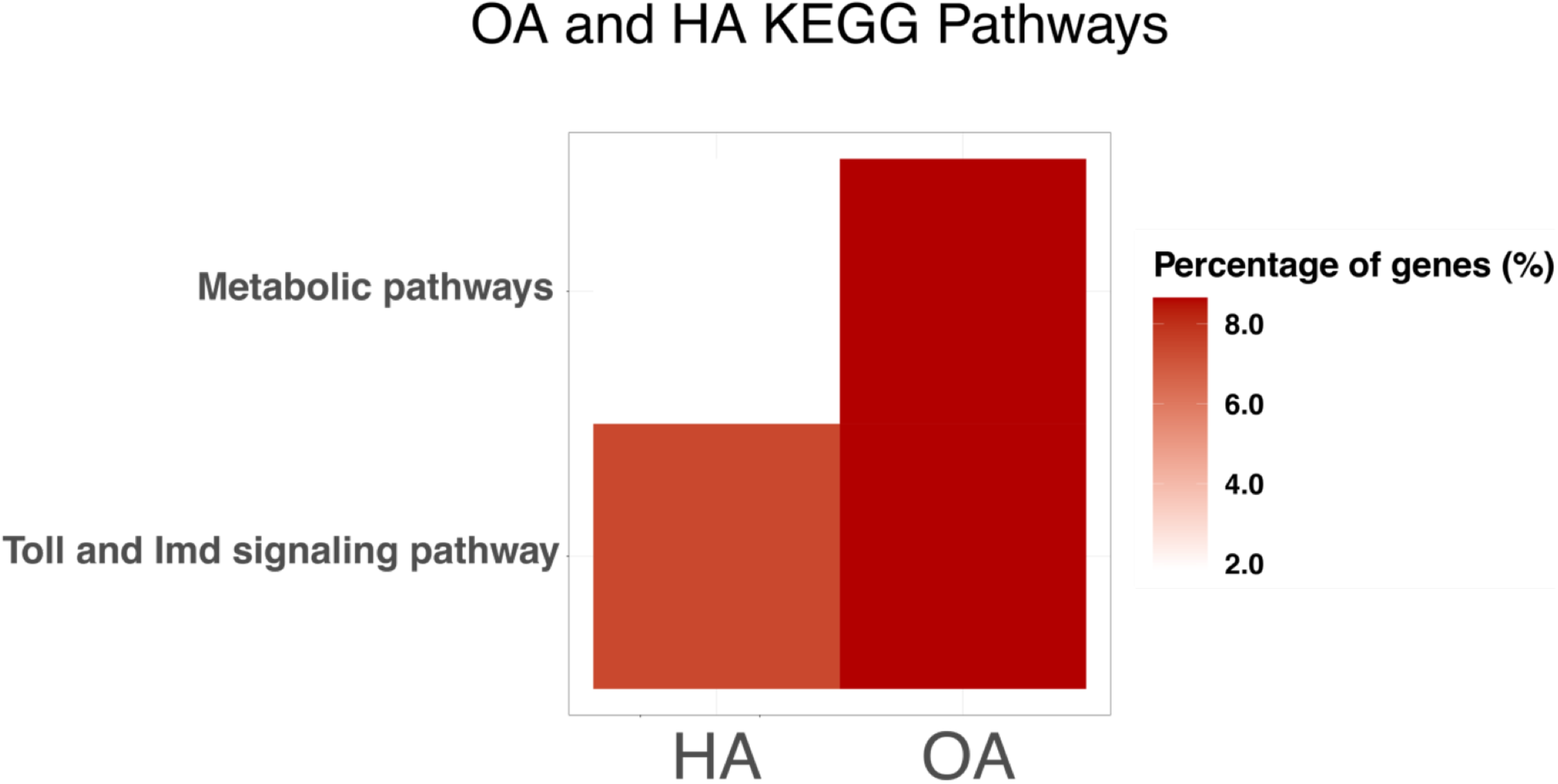
(A) Significantly upregulated genes with *D. melanogaster* orthologs for each species are shown by the percentage of genes of the total number upregulated genes that fall into each KEGG pathway. (B) Significantly downregulated genes with *D. melanogaster* orthologs for each species are shown by the percentage of genes out of the total number of downregulated genes that fall into each KEGG pathway. The KEGG pathways Metabolic pathways and Toll and Imd signaling were excluded. (C) Significantly downregulated genes with *D. melanogaster* orthologs for each species are shown by the percentage of genes out of the total number of downregulated genes that fall into the Metabolic pathways and Toll and Imd signaling pathway KEGG pathways.

### Common transcriptional responses of D. sechellia exposed to HA, OA and L-DOPA suggests overlapping regulatory mechanisms

To assess the overlap in transcriptional responses of *D. sechellia* to both OA and HA, the predominant fatty acid volatiles in *M. citrifolia*, along with 3,4-dihydroxyphenylalanine (L-DOPA), which is found in *M. citrifolia* fruit and is important for the specialization of *D. sechellia* to this fruit (Lanno *et al*. 2019a, Lavista-Llanos *et al*. 2014), we compared DEGs between our HA treatment, DEGs identified in a previous study using a 0.7% OA treatment (Lanno *et al*. 2017), and DEGs identified in a previous study of responses to 10mg/mL of L-DOPA added to the fly food (Lanno *et al*. 2019a). Treatment with OA treatment yielded 103 *D. sechellia* genes with *D. melanogaster* orthologs that were significantly differentially expressed. Treatment with L-DOPA yielded 643 *D. sechellia* genes with *D. melanogaster* orthologs that were significantly differentially expressed. Comparison of genes responsive to HA, OA, and L-DOPA identified 12 DEGs that respond to HA, OA, and L-DOPA treatment in *D. sechellia* (**Figure 3B, Table S6)**. Interestingly, of the 19 shared DEGs with *D. melanogaster* orthologs between OA and HA treatments, all genes are downregulated except for E(spl)mgamma-HLH, which is upregulated upon both OA and HA treatment (**Table S4**). To compare metabolic and cellular pathways involved in the conserved response between OA and HA treatment in *D. sechellia*, GO term enrichment analysis was performed on shared DEGs. Genes involved with biological processes related to the humoral immune response were overrepresented (p = 2.95e-08, **Table S22**). No significantly enriched processes were found for molecular function or cellular component. A KEGG pathway analysis of DEGs in *D. sechellia* similarly found that upon OA, *D. sechellia* downregulated genes involved in both the Toll and Imd signaling pathways as well as genes involved in metabolic processes (**Table S21, Figure S1**). Interestingly, in response to OA and HA, *D. sechellia* downregulated genes involved the humoral immune response (*AttC, CecA2, Def, DptB, Dro, edin*, and *PGRP-SB1* **Table S4**).

## Discussion

Understanding the genetic basis of how organisms evolve to occupy different ecological niches and adapt to their environments is crucial to understanding the evolution of plant and animal interactions. Insect-host plant specialization is an excellent example of the evolution of such interactions and has been the subject of numerous ecological studies. While the phenomenon is well documented, the genetic basis of evolved host specialization is still not widely understood. Here we focus on the specialization of *D. sechellia* to feed almost exclusively on one host plant, *M. citrifolia* because it is an excellent model to understand the genetic basis of dietary specialization. This is in part because it is has evolved recently and very closely related to the genetic model generalist species *D. melanogaster*. Fortuitously, we can take advantage of the wealth of genetic tools and information about *D. melanogaster* and sister species to understand the evolution of dietary specialization in this group (Groen and Whiteman 2016).

*Drosophila sechellia* upregulates a single cytochrome P450 (*Cyp4e1*) in response to HA whereas *D. simulans* downregulates this same gene (**Table S2**), but previous work has shown that cytochrome P450s are not the evolved mechanism by which *D. sechellia* is able to survive OA or HA exposure (Peyser *et al*. 2017). Previous work has shown that HA induces a “reversable coma” in generalist *D. melanogaster, D. simulans*, and *D. mauritiana* flies (Farine *et al*. 1996), and *D. sechellia* prefers to oviposit on HA compared to OA and control food sources (Amlou *et al* 1998). *Drosophila sechellia* has a premature stop codon in *Obp56e* as well as *Obp57d* and *Obp57e* alleles that reduce their avoidance to noni volatiles (Dworkin and Jones 2009, Matsuo *et al*. 2007). The mechanisms that drive this attraction may be through changes in gene expression and may be reflected in these predicted regulatory networks. *Drosophila sechellia* downregulates many genes involved in the Imd and Toll immune pathways when exposed to both HA and OA, suggesting that somehow these two medium chain fatty acids are interacting with negative effectors of Imd signaling. As these two immune pathways have similar and overlapping target genes, further analysis is needed to determine if both pathways are involved in these interactions (Hanson and Lemaitre 2020). This interaction between *D. sechellia* and the volatiles from *M. citrifolia* reduces the immune humoral response, which could cause alterations in gut microbiota composition that aids in the detoxification of relevant plant secondary metabolites. E(spl)mgamma-HLH is a Notch responsive Myc-like transcription factor that has been shown to interact with Relish, the main regulator of Imd immune signaling (Dushay, Åsling, and Hultmark 1996) through recent yeast two-hybrid assays (Shokri *et al*. 2019) and is upregulated in *D. sechellia* in response to both OA and HA exposure (**3B Table S4**). The plastic response of insect immune systems allows them to fend off pathogens when needed, but also allow for the management of endosymbionts (Vilcinskas 2013, Login *et al*. 2011). Unlike its generalist sister species, *Drosophila sechellia* was previously shown to lack an immune response when confronted with parasitic wasps, hinting that immune system responses in this species may be unlike its generalist sister species (Salazar-Jaramillo *et al*. 2017). Additionally, prior work showed that free fatty acids are involved in regulating immune responses in mammals (Alvarez-Curto and Milligan 2016), and hexanoic acid priming of plants can activate the jasmonic acid pathway to increase plant resistance to fungal pathogens (Aranega-Bou *et al*. 2014; García-Robles *et al*. 2013). Further study comparing the microbiomes between *D. sechellia* and *D. simulans* and how they may change from feeding on *M. citrifolia* fruit as well as how each responds to pathogen challenge in this context would help to understand why *D. sechellia* is downregulating its immune response when there could be a serious fitness cost of this action.

Recent studies have shown that *D. melanogaster* uses sweet tasting gustatory receptor neurons to sense OA and HA (Masek and Keene 2013; Tauber *et al*. 2017; Chen and Amrein 2017). Our data shows that upon exposure to HA, only *D. sechellia* significantly upregulates another odorant binding protein, *Obp56a*. In contrast to HA treatment, in response to OA *D. sechellia* adults do not change expression of any odorant binding proteins (Lanno *et al*. 2017). Both *D. simulans* and *D*. melanogaster both downregulate the expression of *Opb57a* in response to HA (**Table S10)**. Recently, studies examining the gustatory and behavioral basis of the attraction of *D. sechellia* to noni have shown that *Or22*a neurons in the fly brain are involved in the attraction to noni (Auer *et al*. 2020), as are *Or85b/c* and *Ir75b* neurons (Prieto-Godino *et al*. 2017). *Orco* mutant *D. sechellia* flies lose olfactory responses to both 2-heptanone and 1-hexanol, two compounds found in ripe noni fruit (Auer *et al*. 2020). Further work on the role of these genes in *D. sechellia* and the regulatory mechanisms responsible for the change of odorant binding protein expression in response to HA may help to elucidate how *D. sechellia* has evolved to specialize on *M. citrifolia*

*Drosophila sechellia* is altering the expression of far fewer genes in response to HA than in the generalist species *D. melanogaster* and *D. simulans* (**Tables S1-3**). *Drosophila sechellia* appears to be downregulating many genes involved in the humoral immune response whereas *D. melanogaster* and *D. simulans* alter the expression of genes involved in many different metabolic pathways (**Figures 4A-4C**). As generalist insects feed on many different plants which produce different secondary metabolites to defend themselves from predators, perhaps generalist species have a more plastic regulatory response to subvert toxicity whereas the specialist *D. sechellia* is resistant to the toxicity of its host through a specific constitutive mechanism. A recent study comparing the fitness of *D. melanogaster* to *D. sechellia* larvae fed different food sources showed a loss of carbohydrate metabolic responses in *D. sechellia*, as they have specialized on a fruit with a relatively low sugar content, *M. citrifolia* (Watanabe *et al*. 2019). Of the many significant genes differentially expressed in *D. melanogaster* and *D. simulans*, most DEGs are not drastically differentially expressed in HA treatment compared to controls (**Figures 2D and 2E**). *Drosophila sechellia* conversely alters the expression of relatively far fewer genes, but many of the DEGs in response to HA are drastically differentially expressed (**Figure 2F**). Similarly, another study found that when adapted to a grass diet, *Spodoptera littoralis* had a smaller transcriptional response when fed maize compared to more generalist *S. littoralis* (Roy *et al*. 2016). Determining the scale of these responsive regulatory effects and their role in toxin resistance will help elucidate how *D. sechellia* has evolved to avoid the toxicity of *M. citrifolia*.

In response to OA, *D. sechellia* increases its expression of several *Osiris* genes (Lanno *et al*. 2017). Previous work using RNAi in *D. melanogaster* to knock-down the expression of individual genes and examine survival in these flies when exposed to OA showed that the reduction of *Osi6, Osi7*, and *Osi8* expression decreased survival (Andrade Lopez *et al*. 2017). In response to HA, neither *D. sechellia* nor *D. melanogaster* significantly alter the expression of any *Osiris* genes, but *D. simulans* downregulates the expression of *Osi6, Osi7*, and *Osi15* (**Table S2**). The cellular and physiological function of these genes is unknown, so understanding what these genes are doing and how they may be helping to shape the interactions between these insects and their toxic hosts may be useful to understand how these interactions evolve (Coolon *et al*. 2019).

Examining and comparing the changes in transcriptional output of insects when exposed to these different plant chemicals in specialized versus generalist species provides a framework to understand how these interactions have evolved. Pathway analyses of these genes is useful in determining the physiological function of altered expression in response to these chemicals, but analyses of transcription factors that alter expression of these genes is necessary to better understand the regulatory mechanisms involved in dietary specialization. Comparing transcription factors responding to plant chemicals may help elucidate regulatory mechanisms involved in these responses and shed light on how insects use changes in the transcription of target genes in order to compete against plants in this evolutionary arms race to adapt to toxic food sources.

## Acknowledgments

Research reported in this publication was supported by Wesleyan University (Startup to JDC, Department of Biology funds to JDC, College of the Environment funds to JDC), and the National Institute Of General Medical Sciences of the National Institutes of Health under Award Number R15GM135901 (award to JDC). The content is solely the responsibility of the authors and does not necessarily represent the official views of the National Institutes of Health.

